# Ancient DNA evidence for the ecological globalisation of cod fishing in medieval and post-medieval Europe

**DOI:** 10.1101/2022.06.03.494519

**Authors:** Lourdes Martínez-García, Giada Ferrari, Angélica Cuevas, Lane M. Atmore, Begoña López-Arias, Mark Culling, Laura Llorente-Rodríguez, Arturo Morales-Muñiz, Eufrasia Roselló-Izquierdo, Juan Antonio Quirós, Ricard Marlasca-Martín, Bernd Hänfling, William F. Hutchinson, Kjetill S. Jakobsen, Sissel Jentoft, David Orton, Bastiaan Star, James H. Barrett

**Affiliations:** Centre for Ecological and Evolutionary Synthesis (CEES), Department of Biosciences, University of Oslo, Oslo, Norway; Royal Botanic Garden Edinburgh, Scotland; Laboratorio de Arqueozoología LAZ-UAM, Universidad Autónoma de Madrid, Madrid, Spain; Evolutionary Biology Group, Department of Biological Sciences, University of Hull, United Kingdom; Laboratory for Archaezoological Studies, Faculty of Archaeology, University of Leiden, Leiden, The Netherlands; Department of Geography, Prehistory and Archaeology, University of the Basque Country, Vitoria-Gasteiz, Spain; Posidonia s.l. Avd. Sant Jordi, Eivissa, Spain; Institute for Biodiversity and Freshwater Conservation, UHI-Inverness, Inverness, United Kingdom; BioArCh, Department of Archaeology, University of York, York, United Kingdom; Department of Archaeology and Cultural History, NTNU University Museum, Trondheim, Norway

**Keywords:** cod trade, historical ecology, marine fisheries, zooarchaeology, biological source

## Abstract

Understanding the historical emergence and growth of long-range fisheries can provide fundamental insights into the timing of ecological impacts and the development of coastal communities during the last millennium. Whole genome sequencing approaches can improve such understanding by determining the origin of archaeological fish specimens that may have been obtained from historic trade or distant water. Here, we used genome-wide data to individually infer the biological source of 37 ancient Atlantic cod specimens (*ca*. 1050 to 1950 CE) from England and Spain. Our findings provide novel genetic evidence that eleventh- to twelfth-century specimens from London were predominantly obtained from nearby populations, while thirteenth- to fourteenth-century specimens derived from distant sources. Our results further suggest that Icelandic cod was exported to London earlier than previously reported. Our observations confirm the chronology and geography of the trans-Atlantic cod trade from Newfoundland to Spain starting by the early sixteenth century. Our findings demonstrate the utility of whole genome sequencing and ancient DNA approaches to describe the globalisation of marine fisheries and increase our understanding regarding the extent of the North-Atlantic fish trade and long-range fisheries in medieval and early modern times.

## 1. Introduction

The expansion of long-range fish trade, not least of Atlantic cod (*Gadus morhua*), has partly driven the development of urbanized market economies across European societies during the last millennium [1-3]. The importance of this trade is well-documented by historical sources from the fourteenth century and can be glimpsed in anecdotal historical records and archaeological evidence from the late eleventh, twelfth and thirteenth centuries [4, 5].

Ancient DNA (aDNA) and stable isotopes have previously shown the early transport of air-dried Arctic Norwegian cod (*stockfish*) to Haithabu in Germany by *ca*. 1066 CE [6, 7]. This exchange developed into a major and wide-ranging Atlantic cod trade across medieval northern Europe, linking towns in Scandinavia, Germany, England, and the Low Countries (e.g., Bergen, Lübeck, King’s Lynn, London, and Deventer) [6-8]. In the Iberian Peninsula, the northern ports were developed as strategic trading posts for receiving and distributing luxury and foreign products from both the Mediterranean and northern Europe [9, 10]. As a consequence, distant-water fisheries and fish trade along the Atlantic coast, from Sevilla to western Ireland and Flanders, started to receive more interest within the Iberian market [11, 12]. Subsequently, post-medieval European expansion to the western Atlantic, especially to Newfoundland, linked the above-mentioned northern and Iberian networks into competing and sometimes complementary long-range fisheries that were sources of both food and wealth [1, 13]. For example, seventeenth-century English catches from Newfoundland were often traded to southern Europe, in an economically significant triangular trade that also entailed salt and wine [14].

Tracing the origin of Atlantic cod specimens harvested for these medieval and post-medieval trade networks contributes to our understanding of economic history and historical ecology. Historical and archaeological sources have revealed the extension of distant-water fisheries and trading networks through time and space [8, 13]. However, the geographical and biological resolution of text-based and archaeological sources is often limited, and the level of detail in such sources often decreases with time depth [15]. Determining the biological origin of archaeological bone assemblages of species such as Atlantic cod can therefore provide important information about the populations targeted through distant-water fishing and/or trade. Since archaeological cod bones can represent local or long-distance (even intercontinental) fishing, it is important to distinguish between source populations. Thus, there has been an increased interest in the use of aDNA and stable isotope methods to identify the origin of archaeological remains to trace the development of the globalisation of marine fisheries [6, 16-20]. Here we use novel whole-genome aDNA approaches to greatly improve the spatial specificity and resolution regarding the inference of source populations of archaeological Atlantic cod bones [21].

We assess the biological origin of 37 Atlantic cod specimens from medieval England (London) and post-medieval Spain (Barcelona, Álava (Castillo de Labastida), Madrid and Sevilla) using low-coverage genome-wide data. We genetically assign such specimens according to patterns of spatial genome-wide differentiation among modern populations of Atlantic cod [22-25]. We specifically investigated significant differentiation in polymorphic chromosomal inversions (i.e., LG1, LG2, LG7, and LG12) [6, 26, 27] that are associated with migratory behaviour and temperature clines [22, 28-31]. Their genetic differentiation can therefore indicate the assignment of specimens towards a particular geographic area [6, 32]. Through these methods, we aim to distinguish source populations with improved discriminating power in relation to previous stable isotope and aDNA approaches [6, 15, 16].

## 2. Materials and Methods

### (a) Sample collection

English samples (*n* = 32) were obtained from eight archaeological locations in London (Figure 1b and Table S1). Based on archaeological evidence and stable isotope analysis, three of the locations (Finsbury Pavement, Seal House and Trig Lane) have previously been inferred to have imported preserved cod [16]. Specimens from the additional five locations (Billingsgate 1982, Cheapside 120, New Fresh Wharf, Nonsuch Palace and Swan Lane) were included to provide a continuous fishing time series from the eleventh to sixteenth-seventeenth centuries CE (Figure 1c). Seven of the archaeological sites in London were urban when occupied. The eighth location (Nonsuch Palace) was a royal residence originally outside London, which was later surrounded by the modern metropolis. Spanish samples (*n* = 5) were obtained from four different archaeological locations: a monastic-upper class context from La Cartuja (Sevilla, late fifteenth-early sixteenth centuries), an urban context from Plaza de Oriente (Madrid, seventeenth-eighteenth century), a context from the fishermen’s quarter from Barraques de pescadors (Barcelona; *ca*. seventeenth century), and a rural castle (Álava) that acted as a military centre during the nineteenth century (J.A. Quirós pers. comm.; Figure 1a and Table S1). Atlantic cod bones are very rare in Iberian archaeological sites [12], therefore, the present five Spanish specimens represent those available for this study.

**Figure 1.**
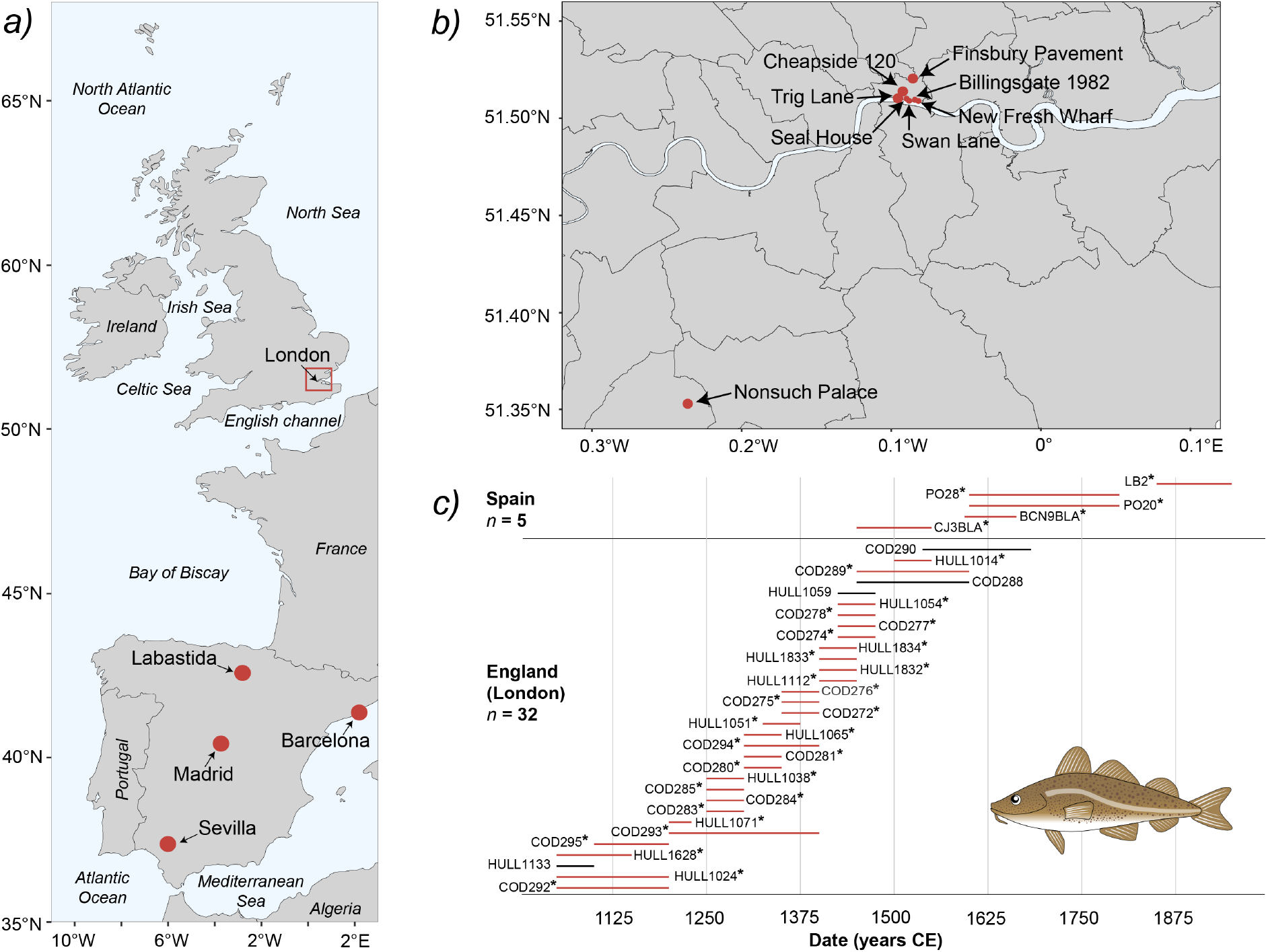
***(a)*** Distribution of archaeological Atlantic cod specimens in England (London) and Spain. Spanish locations are highlighted (in red) on the map. ***(b)*** Detailed distribution (in red) of English archaeological locations in London from which Atlantic cod bones were obtained ***(c)*** Date range of the 37 archaeological specimens as estimated based on archaeological context. Samples in red and with ‘*’ yielded sufficient data to allow more detailed genomic assignments. Other samples could only be assigned to major geographical regions (see results section for explanation). For details regarding the sample codes see Table S1. Fish illustration was drawn by Lourdes Martínez-García.

Cranial (articular, premaxilla, frontal, dentary, and parasphenoid) and postcranial (vertebra and cleithra) bones were included (Table S1 and Figure S2). Cranial bones are more likely to represent local fishing because many preserved fish products were decapitated [33], although, complete fish (and/or preserved fish heads) were sometimes traded over long distances [6, 34, 35]. Cleithra (which support the pectoral fin just behind the cranium) can be found together with cranial remains, or (if fish were decapitated anterior to this element) with postcranial bones [15]. Here, we considered cleithra belonging among the postcranial bones.

After field collection, all samples were stored dry and unfrozen. Dating of the samples was based on archaeological context. Qualitative date ranges were converted into calendar years as per Orton, et al. [15] considering an ‘early’ century the first half of that century (e.g., ‘00 to ‘50), ‘mid’ century as ‘25 to ‘75, and ‘late’ century as the second half of that century (e.g., ‘50 to ‘00). The archaeological Atlantic cod samples were morphologically and genetically identified to species.

### (b) aDNA extraction and library preparation

We processed 18 English (London) fish bone samples in the aDNA laboratory at the University of Oslo [36, 37] (Table S1). Treatment of samples prior to DNA extraction was according to Ferrari, et al. [38] and Martínez-García, et al. [39]. In short, fish bones were UV-treated for 10 minutes per side and milled using a stainless-steel mortar [40]. Milled fish-bone powder was divided in two aliquots per specimen (150-200 mg per aliquot) as starting material for DNA extraction. Genomic DNA was extracted from the fish-bone samples using the mild Bleach treatment and Double-Digestion step (BleDD) protocol [41]. In addition, we added to our initial London assemblage aDNA from 14 English fish bone samples previously processed at the University of Hull following the protocols in Hutchinson, et al. [42] (Table S1). Three out of the 14 samples were previously inferred to have a southern and central North Sea biological origin (Table S1) [42]. Furthermore, we analysed aDNA from five Spanish bone samples previously analysed and processed using a modified protocol of Yang, et al. [43] at BioArch, University of York. In short, samples were decontaminated with 6% bleach for five minutes and then rinsed three times in distilled water. Samples were further UV-treated for 10 to 20 minutes per side. Samples were powdered prior to the addition of a lysis buffer (EDTA) and Proteinase K. Samples were incubated overnight at 50°C while kept in rotation. After incubation, samples were centrifuged to separate bone powder from buffer solution. The supernatant was transferred to an Amicon Ultra-4, Centrifugal Filter Device, 10,000 NMWL tube to concentrate the solution and Quiagen QiaQuick MinElute™ kit was used for DNA purification. Contamination controls were taken during every step of the extraction and amplification procedure.

Double-indexed blunt-end sequencing libraries were built from 16 or 20 μl of DNA extract from all samples using the double-stranded Meyer-Kircher protocol [44, 45] with the modifications listed in Schroeder, et al. [46] or the single-stranded Santa Cruz Reaction (SCR) protocol using tier 4 adapter dilutions [47] (see Table S1 for specifications). Multiple extraction and negative reagent controls during all library sessions were used to detect possible contamination. All samples were assessed for library quality and concentration using a High Sensitivity DNA Assay on the Bioanalyzer 2100 (Agilent) or with a High Sensitivity NGS Fragment Analysis Kit on the Fragment AnalyzerTM (Advanced Analytical). Successful libraries were sequenced using the Illumina HiSeq 4000 with 150 bp paired-reads, or on a Novaseq 6000 with 150 bp paired-reads at the Norwegian Sequencing Centre. Sequencing reads were demultiplexed allowing zero mismatches in the index tag and they were processed using PALEOMIX v1.2.13 [48]. Trimming of residual adapter contamination, filtering and collapsing of reads was done using AdapterRemoval v.2.1.7 [49]. Mapping of remaining reads was performed against the gadMor2 reference genome [50, 51] using BWA v.0.7.12 [52] with the *backtrack* algorithm, disabled seeding and minimum quality score of 25. aDNA deamination patterns were determined using MapDamage v.2.0.9 [53] and BAM files were indexed with samtools v.1.9 [54].

### (c) Genomic and statistical analysis

To infer the biological origin of the archaeological samples of Atlantic cod, we used the BAMscorer pipeline [21] which is especially developed to analyse extremely low-coverage whole genome data. Following the methodology described by Ferrari, et al. [21], we used a genome-wide database of modern individuals that were obtained from Barth, et al. [55] and Pinsky, et al. [24] to further assign ancient specimens. This dataset includes 276 Atlantic cod individuals that represent three broad geographical regions of the species’ range: the western Atlantic Ocean, the eastern Atlantic Ocean and the Baltic Sea. All these regions are genetically differentiated and represent potential sources of distant water fishing and fish trade over the chronology of our study [24, 55]. The genomic assignment to a source population followed a hierarchical procedure (Figure 2a). First, a genome-wide assignment as implemented in BAMscorer [21] (excluding the four large chromosomal inversions in Atlantic cod: LG1, LG2, LG7 and LG12) was used to determine an overall eastern or western Atlantic Ocean origin. Second, all specimens assigned to an eastern Atlantic region were subsequently analysed using a similar approach to determine a possible Baltic Sea origin [21].

**Figure 2.**
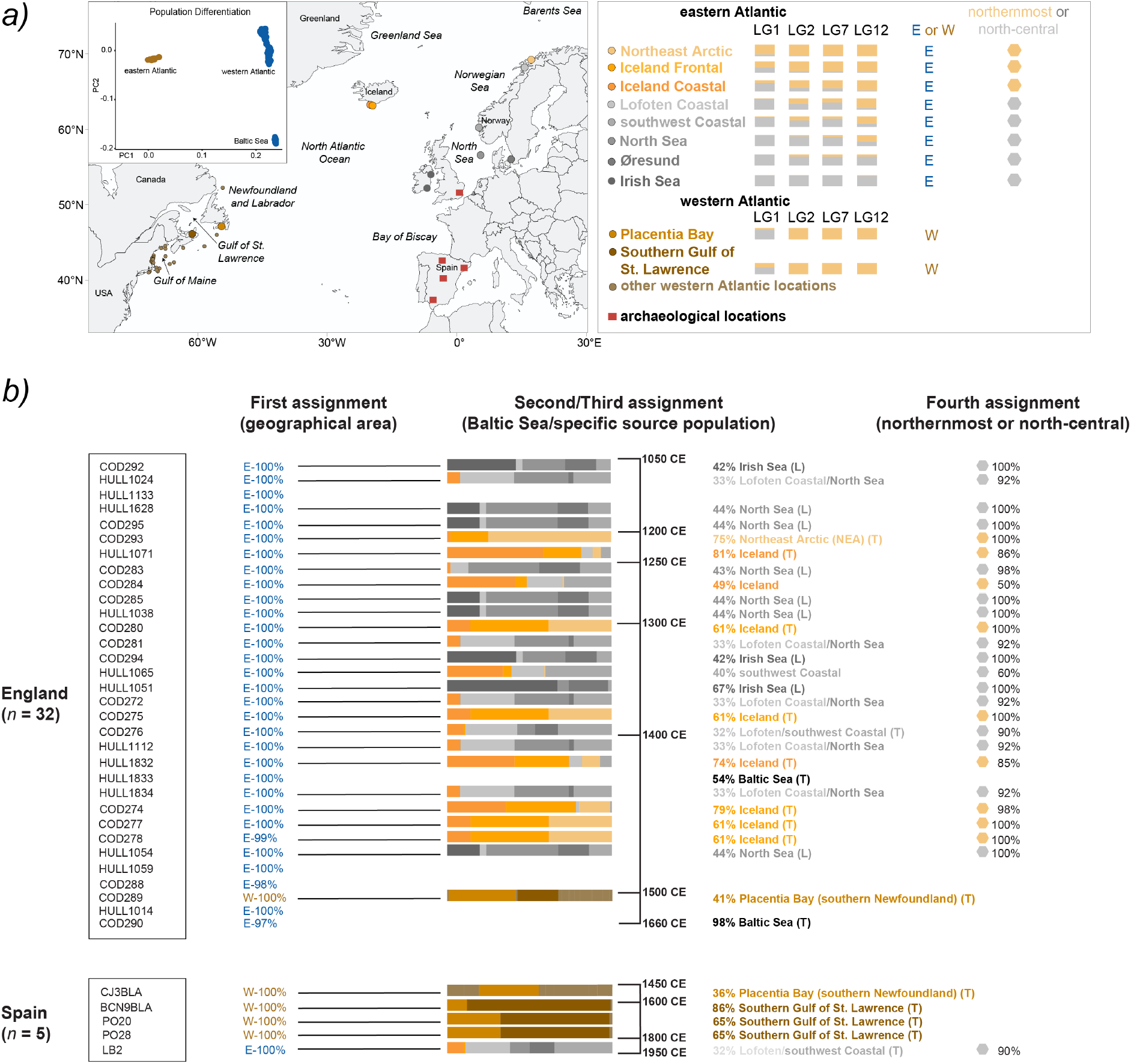
***(a)*** Geographical distribution of inversion frequencies of chromosomal inversions in Atlantic cod (LG1, LG2, LG7 and LG12) across the North Atlantic Ocean. The assignment of specific haplotypes to a geographical area is either eastern (E, in blue) or western (W, in brown) Atlantic. The population PCA plot was modified from Ferrari, et al. [21] and shows the differentiation between the eastern and western Atlantic Oceans, and the Baltic Sea (in blue as it is located within the eastern Atlantic Ocean). Specific alleles associated with a northernmost composite genotype distribution are assigned in orange. Alleles associated with a more temperate north-central genotype distribution are assigned in grey. The archaeological locations are indicated with red squares. The modern populations used as possible source populations of our ancient specimens are located in the map. ***(b)*** First genomic assignment: Overall percentages (%) represent the minimum probability obtained to be from either the eastern or the western Atlantic Ocean. Second/Third genomic assignment: Source population percentages (%) represent the highest probability to be assigned to the Baltic Sea or to a specific modern location (Table S2, S3 and S4). Iceland assignment is obtained by adding the probabilities of both frontal and coastal Icelandic ecotypes. Only two locations from the western Atlantic region have divergent inversion genotype frequencies, thus, they have a specific assignment colour (Placentia Bay and Southern Gulf of St. Lawrence). Other western Atlantic locations are represented in light brown colour (Table S3 and S4). The approximate age (CE) of the specimens is indicated on the right side of the bar plot. Specific time periods are found in Table S1, Figure 1 and Figure S2. A local (L) or traded (T) assignment follows putative source population. Specimens are considered to be of local origin with considerable North Sea or Irish Sea assignments, specimens are considered to have been obtained through trade with significant Northeast Arctic, the Norwegian coast, Iceland, Baltic Sea or western Atlantic assignments. Individuals with ambiguous origin (i.e., HULL1024, COD281, COD272, HULL1112, HULL1834) or with a northernmost or north-central origin below 75% probability (i.e., COD284 and HULL1065) are not identified as local or traded. Individuals COD276 and LB2 are identified as traded specimens as their likely origin is a remote population: Norwegian coast (Lofoten or southwest). Fourth assignment: Percentages (%) representing either the northernmost or north-central genotype distribution after adding the scaled probabilities of selected source populations. For details in the sample codes see Table S1.

Third, we used the individual chromosomal inversion genotypes obtained with BAMscorer [21] to further assign each specimen to a specific source population (Table S2). We used a binomial sampling method as per Star, et al. [6], to infer the probability of the inversion genotype based on spatially divergent and informative inversion frequencies [22-25]. We included those samples where we could infer the genotype of at least three chromosomal inversions with >75% probability (see details in Table S1 and S2). Comparative inversion frequency data were compiled for a range of different populations [22-25]. From the eastern Atlantic we included: the Northeast Arctic, the Norwegian Coast (Lofoten and the southwest), the North Sea, the Irish Sea, Øresund, and Iceland (both coastal and frontal ecotypes, which differ in their migratory behaviour). Western Atlantic populations included a number of populations south and north of Newfoundland (Figure 2a, Table S3 and S4). The highest specific assignment probabilities are reported as percentages (%) representing the confidence with which one individual is assigned to a specific population (i.e., 75%). In the event of similar assignment probabilities for more than one population (e.g., 50% and 50%), both populations are reported as the putative origin of the individual (i.e., Norwegian Coast or North Sea, see Tables S3 and S4 for specific details of assignment percentages). Finally, we recognized that most eastern Atlantic individuals could be further classified with high confidence towards two spatially distinct groups; an overall assignment to a northernmost (Northeast Arctic and Iceland) or north-central (Norwegian Coast, North Sea, Irish Sea and Øresund) distribution (by adding the scaled probabilities of source populations; see details in Table S3 and S4).

We performed a Fisher’s exact test to assess for the existence of an association between bone element and specimen origin (i.e., sourced through trade versus local landings from the North Sea or Irish Sea). The test was implemented in the *stats* and *ggstatsplot* libraries in R [56, 57] using 26 samples that were assigned to a specific source population or the Baltic Sea (Figure 2b and S3). We excluded samples with a northernmost or north-central assignment with less than 75% probability and samples with an indistinguishable origin (see Results section for more details).

## 3. Results

We sequenced 37 specimens and obtained a total of ∼342 million paired reads, ∼9 million aligned reads and endogenous DNA content between <0.01% and 34% (Table S2). Sequencing reads showed the patterns of DNA fragmentation and deamination rates that are consistent with those of authentic aDNA (Figure S1). We successfully assigned these 37 sequenced specimens to one of the two broad geographical areas (eastern or western Atlantic Ocean), finding a total of five samples (England = 1, Spain = 4) from the western Atlantic and 32 specimens (England = 31, Spain = 1) from the eastern Atlantic (Figure 2b, Table S1 and S2). Subsequently, we assigned a total of 33 out of the 37 samples (England = 28, Spain = 5) to either the Baltic Sea or to a more specific source population based on the assignment probabilities of their composite inversion genotypes (Table S2 and Figure 2b). For samples assigned to a specific source population we could identify 111 out of 124 inversion genotypes with more than 95% probability (Table S2). Specific assignments are dependent on the source populations provided for comparison, which can result in low probability assignments as several populations can share similar inversion genotype frequencies (e.g., <50% probability; Figure 2b). Although postcranial bones are more commonly assigned to non-local sources, we did not find a statistically significant association between the bone element (cranial or postcranial) and the origin (local or traded) of the specimen (p = 0.19; Table S1, Figure S2, and S3) with our current sample size.

For London, of the 31 specimens assigned to the eastern Atlantic Ocean, 28 had sufficient data for estimating genotypes of all inversion loci. We found two specimens that could be assigned to the Baltic Sea at 54% and 98% probabilities (Figure 2b). We assigned 16 specimens to a north-central haplotype group (with >60% probability) that includes nine specimens with a possible source population like the North Sea or the Irish Sea (42-67% probability), one specimen with a southwest Norwegian Coast origin (40% probability), and six specimens with indistinguishable associations to the Norwegian Coast (both Lofoten and southwest) and the North Sea (32-33% probability). Nonetheless, these within group assignments are not strongly supported as the inversion frequency differences of the reference populations are limited. Similarly, we assigned eight specimens to a northernmost composite genotype group (with >85% probability), where we found one specimen possibly coming from northern Norway (Northeast Arctic, 75% probability) and seven specimens likely coming from Iceland (61-81% probability). We calculated an overall Icelandic origin by adding the probabilities of being Icelandic frontal or coastal ecotypes. The genomic distinction between Icelandic cod and Northeast Arctic cod is predominantly driven by the higher frequency of north-central genotypes (in grey Figure 2a) for the chromosomal inversion LG01 in Iceland [22]. However, similar inversion frequencies (for LG01) between deep water Icelandic cod and Northeast Arctic cod have been reported [58]. Considering such similarities, the assignments to Iceland or the Northeast Arctic should be taken with caution. Moreover, we found an unreliable assignment for one specimen to a northernmost or north-central composite genotype group (with 50% probability), resulting in an Icelandic assignment with low confidences (49%; Figure 2b, Table S3). Finally, as noted above, we assigned one London specimen (dated between the late fifteenth-sixteenth century) to a western Atlantic origin (with 100% probability) including a possible low confidence assignment to Placentia Bay (41% probability; Figure 2b, Table S3).

For Spain, we found four specimens assigned to the western Atlantic and one specimen assigned to the eastern Atlantic (Figure 2b). The assignments to the western Atlantic (with 100% probability) tentatively included source populations along southwestern Newfoundland (Placentia Bay) or the Gulf of St. Lawrence (with 36-86% probability). We assigned the eastern Atlantic specimen (with 90% probability) to a north-central composite genotype group which includes the Norwegian Coast as a putative origin (indistinguishable association to the Lofoten and southwest Coast with 32-33% probability; Figure 2b, Table S4).

## 4. Discussion

We have used a novel genomic assignment approach to identify the biological source of individual archaeological Atlantic cod specimens from England and Spain. With high confidence, we assigned fish remains to a large-scale geographical origin (up to 100% assignment probability) and genotype groups within regions (>85% assignment probability). With moderate to low confidence (<86% assignment probability), we tentatively identified several of the specimens to more specific spatially constrained populations. Below we describe the resulting time-space patterns observed in England and Spain, and consider the impact these findings have on our understanding of the globalisation of marine fisheries over the last millennium.

### London: an increasingly North Atlantic trade

Earlier zooarchaeological evidence and stable isotope data [15, 16] implied that Atlantic cod trade was predominantly from local fisheries during the eleventh to twelfth centuries, after which longer-distance imports appeared during the thirteenth to fourteenth centuries. Our individual assignments provide confident aDNA evidence that supports this chronology, with fish assigned to northern Atlantic regions appearing in increasing frequency over time. Assignments to specific source populations are often associated with lower individual probabilities (∼32% probability) therefore, these should be considered as indicative only. Interestingly, our assignment analysis indicates that imported Atlantic cod dated between the thirteenth to the fourteenth-centuries derived not only from northern Norway but possibly from Iceland (Figure 2b). According to historical evidence, Iceland first became a major supplier of dried cod to England during the fifteenth century, when fishermen and merchants from England and Germany first defied the Norwegian royal monopoly on trade with Iceland [59, 60]. However, exports of *stockfish* from Iceland via Norway commenced *ca*.1300 CE or earlier [61, 62]. Our results are consistent with this chronology. Icelandic cod in medieval London would probably have reached England via Bergen, on Norwegian, English, and/or Hanseatic ships that are known to have traded between Norway and ports of the English east coast [63, 64].

We also identified samples assigned to the Baltic Sea which is consistent with the beginning of the eastern Baltic cod fisheries by the fifteenth century [17]. Dried cod as a trading product was produced near Gdańsk and Curonia in the Baltic Sea [64]. Furthermore, England’s participation in the western Atlantic cod fisheries expanded in south-eastern Newfoundland during the late sixteenth century (*ca*. 1590; Figure 2b, Table S4) [65]. Therefore, the observation of a specimen from the western Atlantic dated to the late fifteenth to the sixteenth centuries currently represents one of the earliest known genetic examples of this trans-Atlantic expansion. This chronology is consistent with existing knowledge regarding the emergence of trans-Atlantic cod trade, although, as discussed further below, English catches in North America were often destined for southern Europe rather than home markets like London [14, 66].

### Spain: an increasingly trans-Atlantic trade

Like with the single late fifteenth- to sixteenth-century London cod bone, our findings in Spain are consistent with known western Atlantic fishing expansion of the early modern period [67]. In a historical context, the Basques and Galicians provided Atlantic cod for Spain throughout the sixteenth century [66-68]. The fifteenth- to sixteenth-century sample from Spain assigned to waters of the western Atlantic is consistent with historical sources, which indicate that Basque fishermen (from Spain and France) and Galicians acquired fish from southern to western Newfoundland (i.e., Placentia Bay, the Gulf of St. Lawrence, St. Pierre and Miquelon) to fulfil the demand for Atlantic cod in Spain [67, 69-72]. The three later specimens (seventeenth century and seventeenth to eighteenth centuries) could have derived from Spanish fishermen operating in Newfoundland [68, 72] or as trade items with English or French fishermen that had been operating in these grounds since the sixteenth century. In fact, the English engaged in unofficial trade even when political hostilities disrupted relations with Spain [66]. These specimens may thus relate to the triangular trade involving an exchange of Atlantic cod from the western North Atlantic for wine and salt from southern Europe [14]. By the late nineteenth to early twentieth century, dried Atlantic cod from across the Norwegian waters could have been used to provision the military centre in Álava (J.A. Quirós pers. comm.). Our most recent Spanish sample originated from the Norwegian Coast (Figure 2b and Table S4), is therefore consistent with the supply of northern European air-dried Atlantic cod since the eighteenth century [73, 74].

## Conclusion

Altogether, our results provide genetic evidence for an expanding trade and increasing demand for marine fish leading to the exploitation of a great diversity of distant-water sources already in the Middle Ages. Our evidence also tracks the culmination of the marine fisheries extension with European exploitation of the western Atlantic fishing grounds around Newfoundland, starting in the sixteenth century. Our findings emphasise the utility of whole-genome sequencing and ancient DNA methods to describe the increasing demand for Atlantic cod for European societies during medieval and postmedieval periods. These results corroborate and significantly increase existing knowledge about the globalisation of marine fisheries and fish trade in medieval and early modern times.

## Data accessibility

The raw reads for the ancient specimens are released under the ENA accession number PRJEB52865.

## Author contributions

Conceptualization and project design: JHB, BS and LMG. Laboratory work: LMG, GF, AC, LMA, MC, LLR and BLA. Genomic data curation: LMG and GF. Statistical analysis: LMG. Ancient Atlantic cod specimens and archaeological context information: LLR, AMM, ERI, JAQ, RMM, DO and JHB. Resources: LLR, AMM, ERI, DO, BH, WFH and JHB. Data visualization: LMG and BS. Supervision: JHB and BS. Funding acquisition: BS, KSJ, SJ, WFH, DO and JHB. Original draft: LMG. Review & editing: All authors.

## Acknowledgements

We thank Lori Lawson Handley for providing ancient DNA resources for this research. We thank Martin Biddle (Oxford University), Francis Grew (Museum of London), Cath Maloney (Museum of London), Natasha Powers (MOLA) and Roy Stephenson (Museum of London) for assistance with sampling and sampling permissions. We thank M. Skage, S. Kollias, A. Tooming-Klunderud and the Ullevål Sequencing team at the Norwegian Sequencing Centre for sequencing and processing of the genomic samples. Analyses were performed on resources provided by UNINETT Sigma2 – the National Infrastructure for High Performance Computing and Data Storage in Norway.

