## Supplementary_material for "Ancient DNA evidence for the ecological globalisation of cod fishing in medieval and post-medieval Europe"

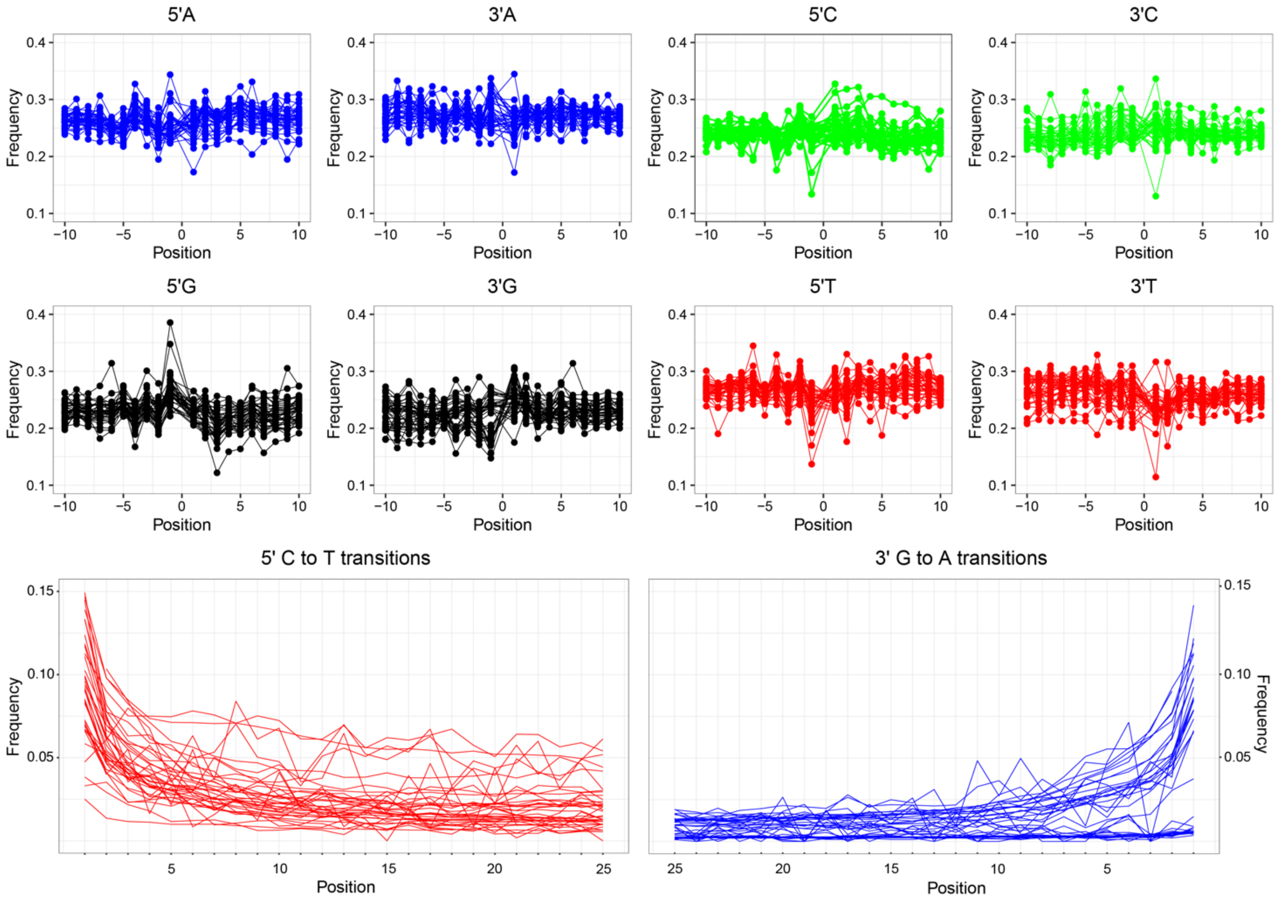


**Figure S1.** Typical fragmentation and misincorporation patterns of nucleotides of aDNA from sequencing data of 37 Atlantic cod specimens from England (London) and Spain. At the top, we show base frequencies. At the bottom, we show the increase in cytosine to thymine (C > T) misincorporations due to cytosine deamination at the 5′-end of DNA fragments and the corresponding increase of guanine to adenine (G > A) misincorporations at the 3′-end. Low quality specimens (e.g., low number of reads) introduce undefined patterns in the Cytosine to thymine (C > T) misincorporations.

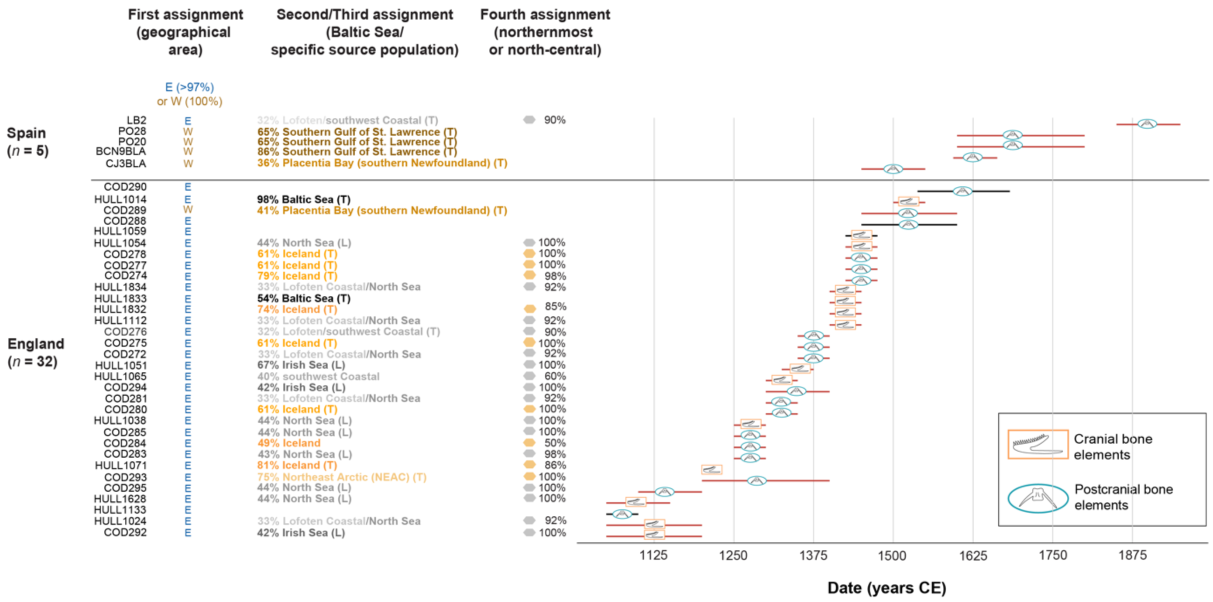


**Figure S2.** Chronological cranial and postcranial bone element distribution of 37 Atlantic cod specimens. First genomic assignment: eastern (E- in blue) or western (W-in brown) Atlantic. Second/Third genomic assignment: assignment to the Baltic Sea or to a specific source population. Iceland assignment is obtained by adding the probabilities of both frontal and coastal Icelandic ecotypes. In parenthesis L indicates specimens with a likely local origin (from the North Sea or Irish Sea), while T indicates possible traded specimens (from other localities). Individuals with ambiguous origin and with a northernmost or north-central origin below 75% probability are not identified as local or traded (see details in Figure S3). Fourth genomic assignment: northernmost (in orange) or north-central (in grey) group. For more information about genomic assignments see methods section in the main text. Dentary (cranial bone element) and vertebra (postcranial bone element) illustrations were drawn by Lourdes Martínez-García.

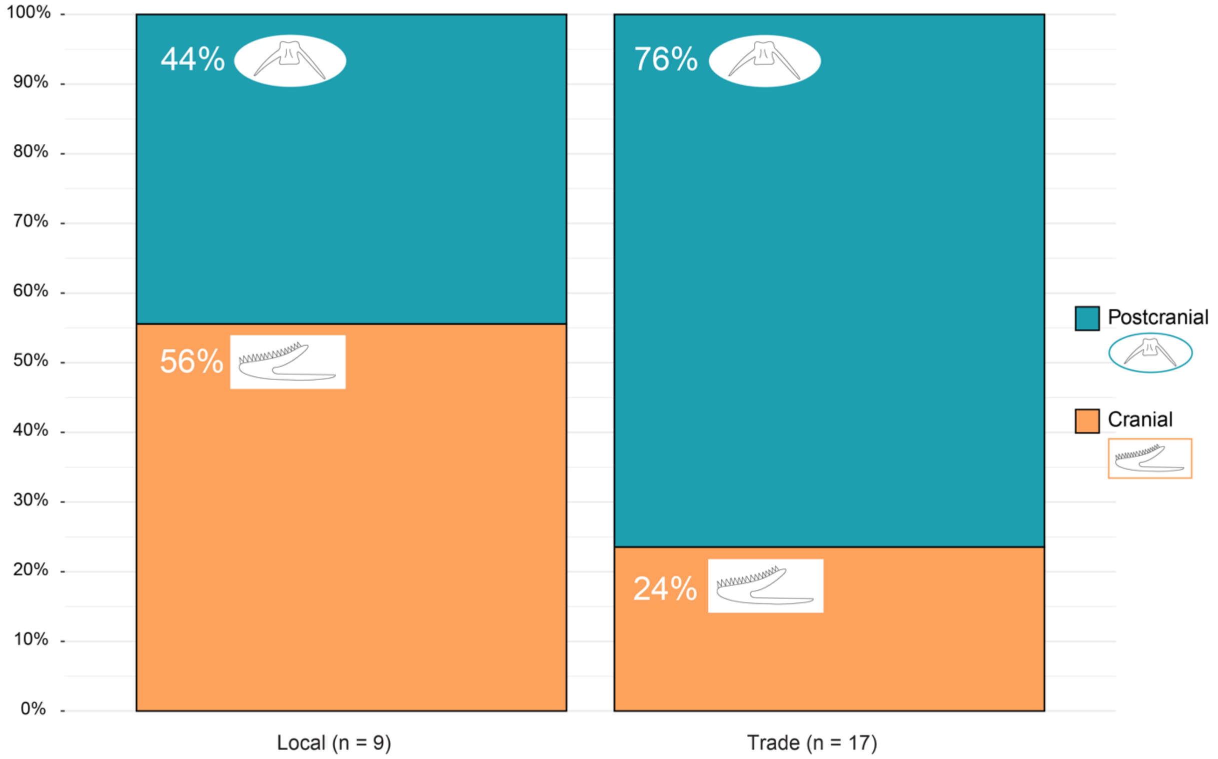


**Figure S3.** Percentage (%) of cranial (orange) or postcranial (blue) bones with a local (North Sea or Irish Sea) or traded (rest of populations) origin. Individuals with a confident assignment to a source population or the Baltic Sea ($n$ = 26; see Table S3 and S4) were included for the Fisher's exact test ($p$ = 0.19). Individuals with ambiguous origin (i.e., HULL1024, COD281, COD272, HULL1112, HULL1834) and with a northernmost or north-central origin below 75% probability (i.e., COD284 and HULL1065) were excluded from this test. Individuals COD276 and LB2 were included in the test as their likely origin is a traded population: Norwegian coast (Lofoten or southwest). Dentary (cranial bone element) and vertebra (postcranial bone element) illustrations were drawn by Lourdes Martínez-García.
